# Defining the Ligand-dependent Interactome of the Sigma 1 Receptor

**DOI:** 10.1101/2022.06.22.497210

**Authors:** Jing Zhao, Rajalakshmi Veeranan-Karmegam, Frederick C. Baker, Barbara A. Mysona, Pritha Bagchi, Yutao Liu, Sylvia B. Smith, Graydon B. Gonsalvez, Kathryn E. Bollinger

## Abstract

Sigma 1 Receptor (S1R) is a therapeutic target for a wide spectrum of pathological conditions ranging from neurodegenerative diseases to cancer and COVID-19. S1R is ubiquitously expressed throughout the visceral organs, nervous, immune and cardiovascular systems. It is proposed to function as a ligand-dependent molecular chaperone that modulates multiple intracellular signaling pathways. The purpose of this study was to define the S1R interactome under native conditions and upon binding to well-characterized ligands. This was accomplished by fusing the biotin ligase, Apex2, to the C terminus of S1R. Cells stably expressing S1R-Apex or a GFP-Apex control were used to map specific protein interactions. Biotinylated proteins were labeled under native conditions and in a ligand dependent manner, then purified and identified using quantitative mass spectrometry. Under native conditions, S1R biotinylates over 200 novel proteins, many of which localize within the endomembrane system (ER, Golgi, secretory vesicles) and function within the secretory pathway. Under conditions of cellular exposure to either S1R agonist or antagonist, results show enrichment of proteins integral to secretion, extracellular matrix formation, and cholesterol biosynthesis. Notably, Proprotein Convertase Subtilisin/Kexin Type 9 (PCSK9) displays increased binding to S1R under conditions of treatment with Haloperidol, a well-known S1R antagonist; whereas Low density lipoprotein receptor (LDLR) binds more efficiently to S1R upon treatment with (+)-Pentazocine ((+)-PTZ), a classical S1R agonist. Our results are consistent with the postulated role of S1R as an intracellular chaperone and further suggest important and novel functionalities related to cholesterol metabolism and biosynthesis.

## INTRODUCTION

The sigma 1 receptor (S1R) is a small (25kD), ubiquitously expressed, transmembrane protein that is localized within the endoplasmic reticulum and its mitochondria-associated membranes(1,2). Studies also show that it can translocate to nuclear and plasma membranes under certain conditions(3-5). S1R has been the subject of intense pharmacologic analysis over the past several decades due to the therapeutic potential of its ligands. In many instances, S1R-mediated therapeutics are advancing from bench to bedside. For example, clinical trials targeting S1R for treatment of neurodegenerative diseases including Amyotrophic Lateral Sclerosis (ALS) and Huntingtons Disease are ongoing(6). In addition, promising preclinical studies indicate that S1R may offer a treatment target for visual disorders including glaucoma, retinal degeneration and traumatic optic neuropathy(7-12). Furthermore, in recent studies, S1R was shown to link the SARS-CoV2 replicase/transcriptase complex to the ER membrane by binding directly to nonstructural protein 6 (NSP 6)(13). Therefore, S1R ligands may provide antiviral activity against severe acute respiratory syndrome CoV-2 (SARS-CoV-2)(13,14). Finally, there is interest in using S1R ligands for treating and imaging cancer(15,16).

Despite intense interest, the molecular mechanisms that underlie effects of S1R ligands are not well understood. In general, agonists for S1R show pro-survival effects whereas antagonists inhibit tumor cell proliferation and induce apoptosis(17). Studies indicate that S1R functions as a “pluripotent modulator”of multiple signaling pathways and therefore affects a wide range of cellular activities including calcium homeostasis, ion channel regulation, and responses to ER and oxidative stress(18). The proposed general mechanism for S1R function is through protein-protein interactions(1). In support of this paradigm, S1R has been reported to bind to at least 49 different proteins that generally show diverse structure and function(19). However, published experiments used to support direct interactions between S1R and other proteins have been mainly limited to low-throughput, candidate-based methods. These include co-immunoprecipitation and resonance energy transfer experiments(5,20,21). Previous studies have also utilized proximity ligation assays, but have not combined these evaluations with high-throughput proteomic analyses.

In this study, a cell line-based proximity biotin labeling assay was developed and combined with proteomic identification(22). The promiscuous biotin ligase Apex2 fused to the C-terminus of S1R was used to label interacting proteins under native conditions and in a ligand dependent manner(23,24). The biotinylated proteins were then purified and identified using quantitative mass spectrometry. Under native conditions, we find that S1R interacts with over 200 novel proteins, many of which localize within the endomembrane system (ER, Golgi, secretory vesicles) and function within the cellular secretory apparatus. In addition, under conditions of cellular exposure to either S1R agonist or antagonist we identify interactome changes that highlight the molecular pathways critical to S1R-mediated ligand-dependent effects. These include proteins involved in cholesterol and lipid metabolism as well as matricellular and extracellular proteins. Our results offer high-throughput evidence that S1R is indeed capable of interacting with a vast array of proteins, consistent with its postulated role as an intracellular chaperone.

## RESULTS

### Defining the interactome of the Sigma1 receptor

In order to identify interacting partners of the Sigma1 receptor (S1R), we employed a proximity biotinylation approach. With this strategy, the protein of interest is tagged with a promiscuous biotin ligase. Proteins that are present within 15nm of the tagged protein become biotinylated in vivo and can be easily purified using streptavidin conjugated beads (Fig. 1A)(25). One of the main advantages of this approach is that because the biotin-streptavidin interaction is extremely strong, the purification can be done using harsh wash buffers. This minimizes binding of non-specific proteins to beads, thus reducing the potential for false-positive hits(26). For our studies, we chose to tag S1R at the C-terminus with Apex2 (27-29). We chose this approach because previous studies have shown that S1R-Apex is functional and produces the expected localization pattern(27). A V5 epitope was incorporated into the tag in order to enable western blotting and immunofluorescence analysis using commercially available antibodies. A construct expressing GFP-Apex was used as the negative control.

**Figure 1:**
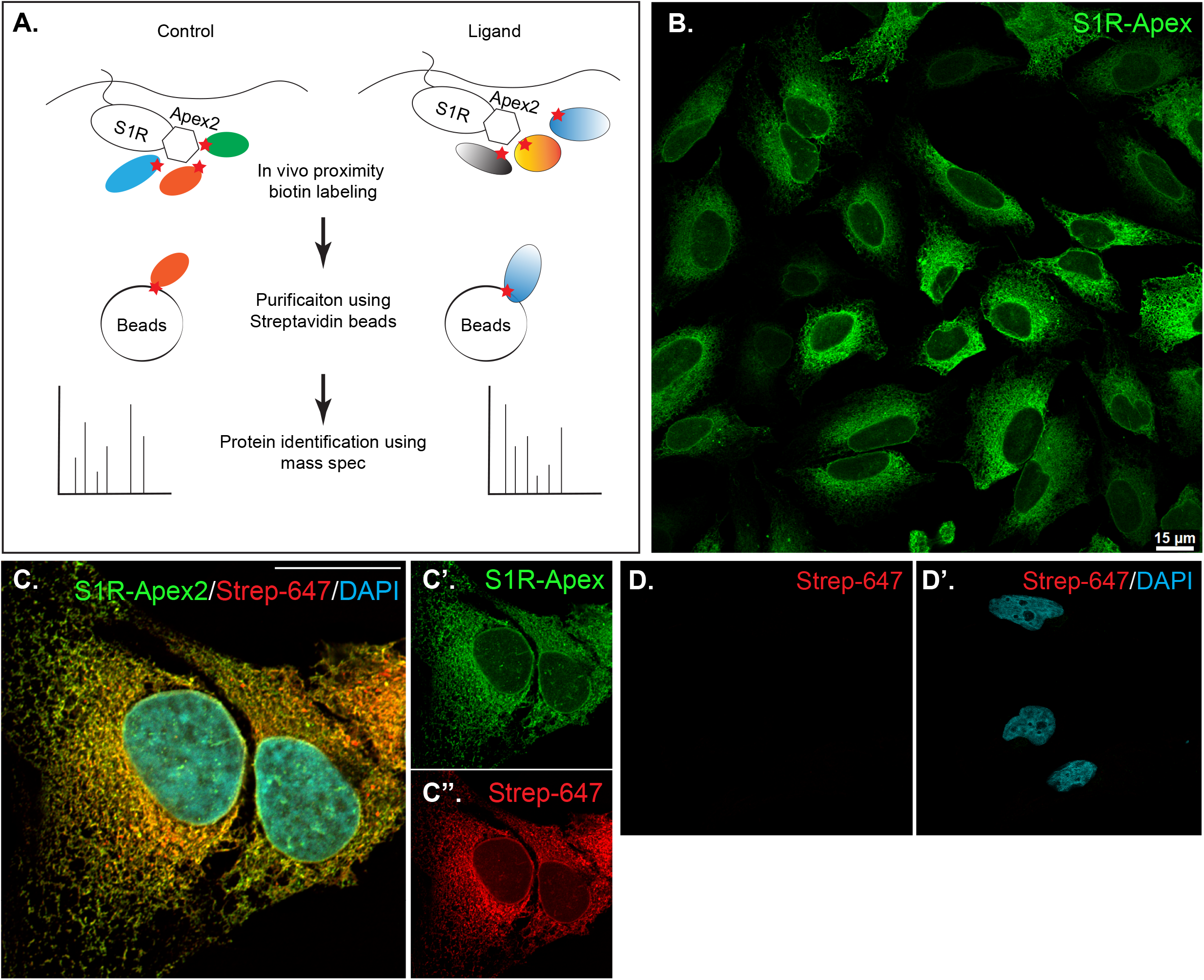
Generation and validation of S1R-Apex cells. **A)** Schematic of the proximity biotinylation strategy **B)** HeLa cells stably expressing S1R-Apex were fixed and processed for immunofluorescence using a V5 antibody (green). S1R-Apex localizes around the nuclear envelope and to ER tubules. **C)** HeLa cells stably expressing S1R-Apex were fixed and processed for immunofluorescence using a V5 antibody (green). The cells were also incubated with Streptavidin-647 (red) to reveal the localization of biotinylated proteins and were counterstained using DAPI (cyan). Biotinylated proteins co-localize with S1R-Apex. **D)** Control HeLa cells not expressing a biotin ligase were processed using Streptavidin-647 (red). The cells were counterstained with DAPI (cyan). Minimal Streptavidin signal is observed in these cells. The scale bar in B is 15 microns and the scale bar in C is 20 microns.

HeLa cells stably expressing S1R-Apex and GFP-Apex were generated using CRISPR-based incorporation of the constructs at the *AAVS1* safe harbor locus using a published protocol(30). Immunostaining of cells expressing S1R-Apex produced the expected localization pattern (Fig. 1B). In order to validate the proximity biotinylation approach, cells expressing S1R-Apex were labeled using biotin phenol. Control cells that did not express a biotin ligase were similarly treated. The cells were then processed for immunofluorescence using a V5 antibody for detecting the Apex tagged protein, and Streptavidin conjugated Alexa647 for detecting biotinylated proteins. In contrast to control cells, robust biotinylation signal was observed in cells expressing S1R-Apex (Fig. 1C, D). Furthermore, the biotinylation signal co-localized with the V5 signal for S1R-Apex (Fig. 1C). The biotinylation pattern is consistent with the published localization of S1R to ER and nuclear membranes (4,27,31). By contrast, cells expressing GFP-Apex displayed a nuclear and cytoplasmic biotinylation pattern (Supplemental Fig. 1). Based on these results, we conclude that S1R-Apex is localizing as expected and biotinylating proximal interacting proteins.

Next, cells expressing GFP-Apex and S1R-Apex were grown and labeled for proteomics analysis. Lysates were prepared from the labeled cells and the biotinylated proteins were purified using high-capacity Streptavidin agarose beads. After extensive wash steps, the bound proteins were eluted using Trypsin digestion and processed for mass-spectrometry. The entire experiment was done in triplicate. Proteins that were enriched at least two-fold in the S1R-Apex pellet in comparison to GFP-Apex and had a p value of at least 0.05 were considered to be specific interacting partners (Fig. 2A, Supplemental Table 1). Using these criteria, S1R interacts with 233 specific proteins in HeLa cells. The top 60 S1R interacting candidates are shown in Table 1.

**Figure 2:**
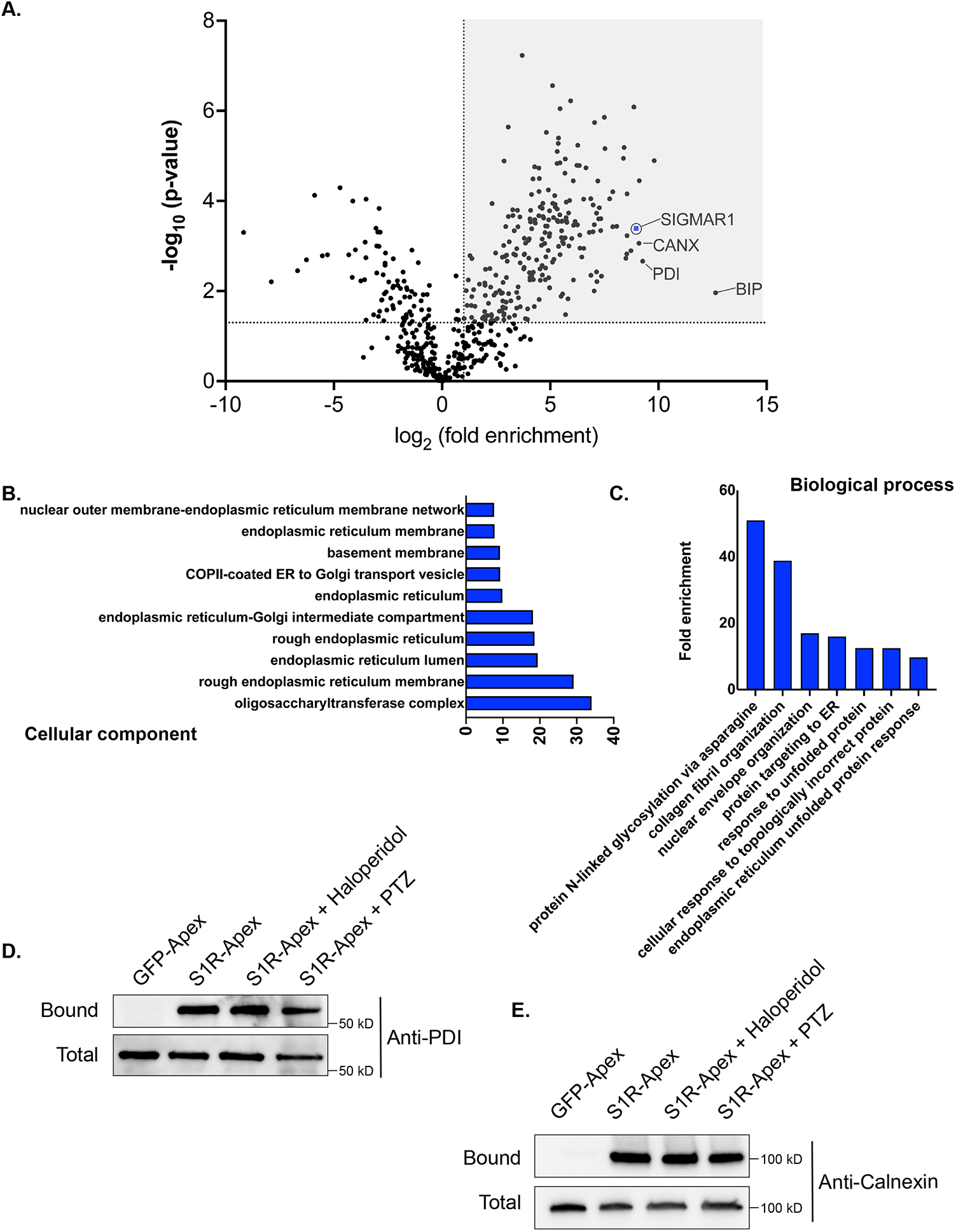
The S1R interactome. **A)** A volcano plot depicting the S1R interactome. A line demarcating 2-fold enrichment and a p value of 0.05 is shown. The grey shaded box indicates proteins that are at least 2-fold enriched with S1R in comparison to the GFP-Apex control with a p value of at least 0.05. **B)** A Cellular Component GO analysis of S1R interacting proteins. **C)** A Biological Process GO analysis of S1R interacting proteins. **D-E)** Biotinylated proteins were purified from HEK293T cells stably expressing either GFP-Apex or S1R-Apex using streptavidin conjugated beads. Biotinylated proteins were also purified from HEK293T cells stably expressing S1R-Apex that were treated with either Haloperidol or (+)-PTZ. The bound proteins were eluted and analyzed by western blotting using an antibody against PDI (D) or Calnexin (E). S1R expressed in HEK293T cells is able to interact with PDI and Calnexin. This interaction is not altered by treatment with Haloperidol or (+)-PTZ.

S1R is known to exist as a trimeric complex in vivo(28). Thus, S1R-Apex should interact with endogenous S1R and result in its biotinylation and enrichment within the dataset. Consistent with this notion, endogenous S1R was highly enriched in the S1R-Apex pellet (Fig. 2A). In addition, the ER chaperone BiP (GRP78), a known S1R interacting partner, was also specifically enriched in the S1R-Apex pellet (Fig. 2A)(1). Previous studies have shown that S1R localizes to the endoplasmic reticulum (ER), to nuclear lamellae, to sites of ER-mitochondria contact and to the plasma membrane(1,3,5,27). Consistent with these studies, a “cellular component” Gene Ontology (GO) analysis indicates that the S1R interactome is highly enriched for proteins that localize to the nuclear membrane, to the ER lumen and membrane, and to organelles involved in protein secretion. (Fig. 2B). Previous studies have implicated a role for S1R in the ER stress response and as a potential molecular chaperone. In line with these findings, a “biological process” GO analysis is enriched for terms such as “protein targeting to ER” and “response to unfolded protein”. In addition, proteins with a role in N-linked glycosylation are highly enriched within the S1R interactome (Fig. 2C).

In order to validate these results, we generated HEK293T cells that stably express either GFP-Apex or S1R-Apex integrated at the *AAVS1* safe harbor locus (30). Two ER proteins that were highly enriched in the S1R interactome were Protein Disulfide Isomerase (PDI) and the chaperone Calnexin. Consistent with the proteomics results, PDI and Calnexin were specifically biotinylated and precipitated in HEK293T cells expressing S1R-Apex (Fig. 2D, E). The interactions between S1R and these proteins were unchanged upon treating cells with either Haloperidol, an S1R antagonist, or with (+)-PTZ, an S1R agonist (Fig. 2D, E).

### S1R interacts with the ER translocation machinery

For proteins that contain a signal sequence, import into the ER occurs co-translationally. Once the signal sequence has been translated, it is bound by the signal recognition particle (SRP). This complex is then docked onto the surface of the ER by binding to the SRP receptor. Next, the docked ribosome associates with the Sec61 translocation complex and its accessory proteins. This enables the translating peptide to be imported into the ER. Once in the ER, luminal chaperones such as BiP and Calnexin help fold the protein into its native conformation (Fig. 3A)(32,33). Multi-pass transmembrane proteins are imported into the ER with the aid of the Nicalin-TMEM147-NOMO complex(34). Numerous proteins within the S1R interactome function at various steps in the ER translocation process (Fig. 3B). This finding was quite intriguing given the proposed function of S1R as a molecular chaperone. These results position S1R at the entry way into the ER, an ideal location for a molecular chaperone.

**Figure 3:**
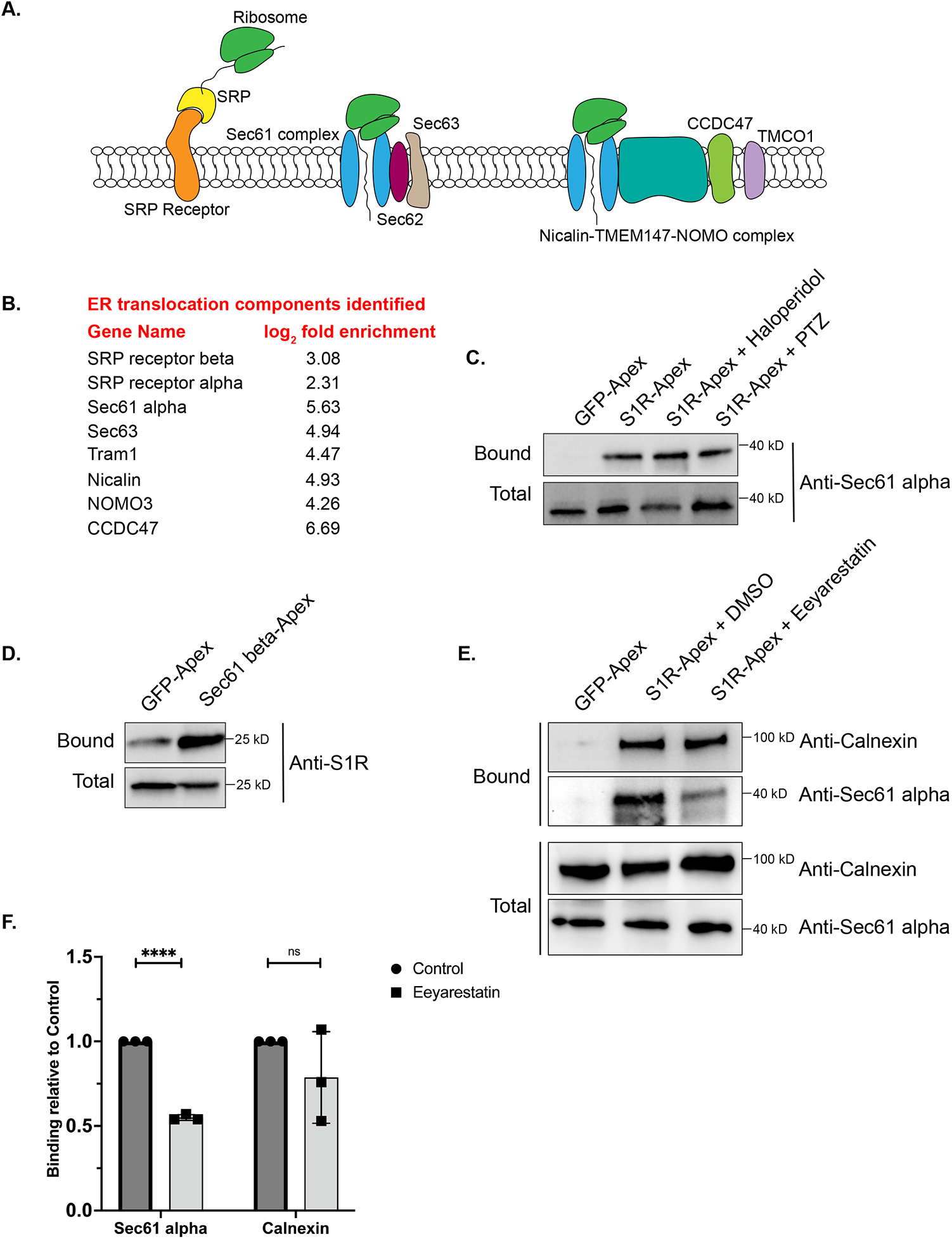
S1R interacts with components of the ER translocation complex. **A)** A schematic of the components involved in translocation of proteins into the ER. **B)** A list of ER translocation components identified in the S1R interactome **C)** Biotinylated proteins were purified from HEK293T cells stably expressing either GFP-Apex or S1R-Apex that were untreated or treated with either Haloperidol or (+)-PTZ. The proteins were purified using streptavidin conjugated beads. The bound proteins were eluted and analyzed by western blotting using an antibody against Sec61alpha. S1R expressed in HEK293T cells interacs with Sec61alpha. **(D)** HEK293T cells were transiently transfected with a construct expressing either GFP-Apex or Sec61 beta-Apex. Biotinylated proteins were purified using streptavidin conjugated beads, the bound proteins were eluted and analyzed by western blotting using an antibody against endogenous S1R. Endogenous S1R interacts with Sec61 beta, a component of the ER translocation complex. **(E)** HEK293T cells stably expressing GFP or S1R-Apex were either untreated (DMSO) or treated with the drug Eeyarestatin. Eeyarestatin blocks the transport of proteins across the Sec61 translocation channel. Biotinylated proteins were purified using streptavidin conjugated beads and the bound proteins were eluted and analyzed by western blotting using antibodies against Sec61 alpha and Calnexin. **(F)** The above experiment was repeated in triplicate and the binding of S1R-Apex with Sec61 alpha and Calnexin was quantified. Unpaired *t* tests were used for these analyses; ****p<0.0001, ns = not significant. Blocking protein transport across the Sec61 channel disrupts the interaction between S1R and Sec61 alpha.

In order to validate these findings, we focused our studies on Sec61alpha, the protein that forms the central translocation channel. Consistent with the proteomics result, Sec61alpha was specifically detected in the S1R-Apex pellet from HEK293T cells (Fig. 3C). As with PDI and Calnexin, the interaction between S1R and Sec61alpha remained unchanged upon treating cells with either Haloperidol or (+)-PTZ. As further validation of this result, we determined whether endogenous untagged S1R was also present in a complex with the ER translocation channel. For this experiment, HEK293T cells were transiently transfected with either GFP-Apex or Sec61beta-Apex. After binding and wash steps, the bound proteins were eluted and analyzed by western blotting using an antibody against S1R. Significantly more S1R was found to be in a complex with Sec61beta-Apex in comparison to GFP-Apex (Fig. 3D), thus validating the interaction between S1R and the ER translocation complex.

We next determined whether the interaction between S1R and the Sec61 complex was sensitive to active protein import into the ER. The drug Eeyarestatin is known to inhibit Sec61 alpha, and as a consequence, ER protein import is blocked (35-37). Interestingly treating cells with Eeyarestatin reduced the interaction between S1R and Sec61alpha (Fig. 3E, F). The interaction between S1R and Calnexin was also somewhat reduced. However, these results were not statistically significant (Fig. 3E, F). Based on these results, we conclude that S1R interacts with the ER translocation machinery and likely does so in an import-dependent manner.

### S1R interacts with components at the ER-Golgi intermediate compartment

Another category of proteins enriched in the S1R interactome are proteins that localize to the ER-Golgi intermediate compartment (ERGIC) (Fig. 2B). Proteins that localize at this site are involved in trafficking between the ER and Golgi compartments and often have a role in the secretory process(38). The S1R interactome contains several proteins that are secreted or localized on the plasma membrane such as Collagens, Integrins, and Low Density Lipoprotein Receptor (Supplemental Table 1). It is therefore possible that S1R plays an active role in the secretory process by interacting with components of the ERGIC. In order to validate this finding, the binding experiment was repeated, and the pellets were analyzed by western blotting using an antibody against Lman1, a marker protein of the ERGIC and a protein that was highly enriched in the S1R proteomics dataset (Supplemental Table 1)(39). Lman1 specifically interacts with S1R-Apex and this interaction is unchanged upon treating cells with either Haloperidol or (+)-PTZ (Fig. 4A). In addition to this biochemical interaction, we observed significant co-localization between S1R and GFP-Lman1 (Fig. 4B).

**Figure 4:**
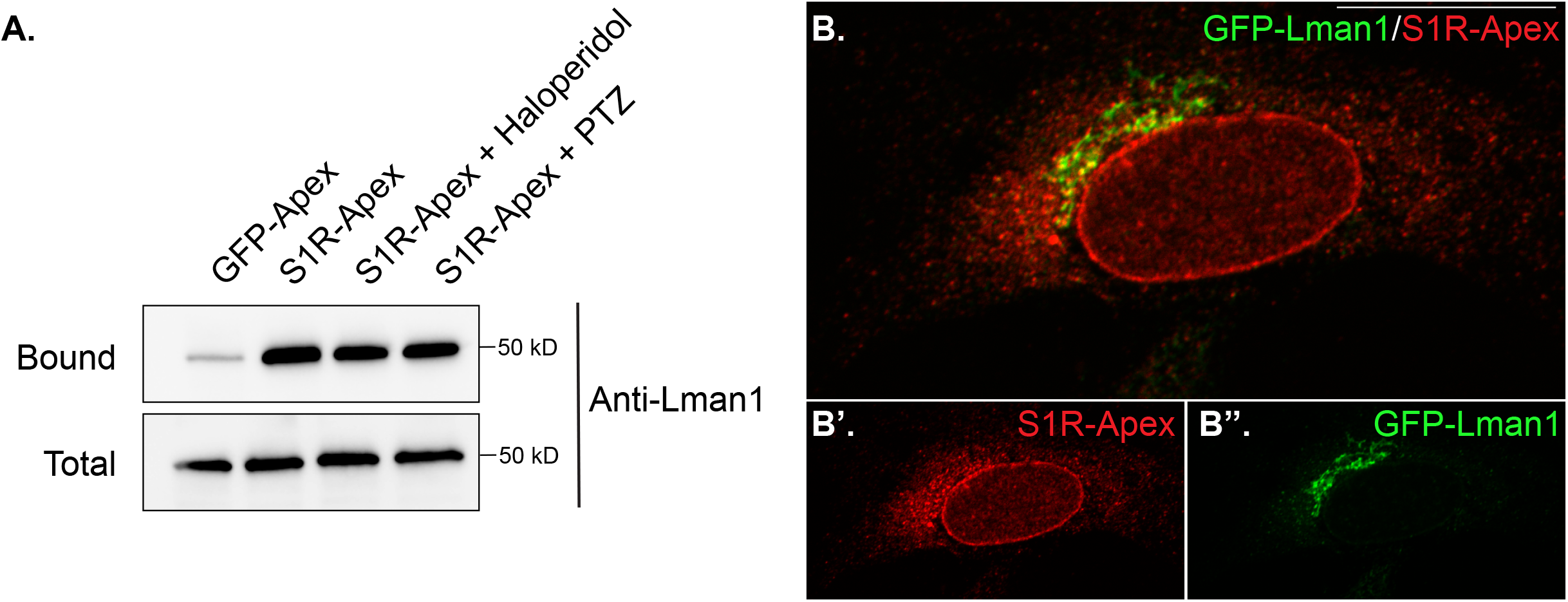
S1R interacts with the ER-Golgi intermediate compartment component Lman1. **(A)** Biotinylated proteins were purified using streptavidin conjugated beads from HEK293T cells stably expressing either GFP-Apex or S1R-Apex that were untreated or treated with either Haloperidol or (+)-PTZ. The bound proteins were eluted and analyzed by western blotting using an antibody against Lman1. S1R expressed in HEK293T cells interacs with the ERGIC component Lman1. **(B)** HeLa cells stably expressing S1R-Apex were transfected with a plasmid expressing GFP-Lamn1 (green). The cells were fixed and processed for immunofluorescence using a V5 antibody (red). S1R-Apex co-localizes with GFP-Lman1 in the area of the ER-Golgi intermediate compartment.

### Ligand dependent interactome of S1R

Having established the S1R interactome under native conditions, we next wished to determine how the interactome would change when cells were treated with Haloperidol, an S1R antagonist, or with (+)-PTZ, an agonist of S1R (40-42). For this experiment, HeLa cells expressing S1R-Apex were either untreated, treated with 25uM Haloperidol for 24 hours, or treated with 20uM (+)-PTZ for 24 hours. The treated cells were labeled, the biotinylated proteins were purified using streptavidin agarose and the bound proteins were analyzed using mass-spectrometry. As before, the entire experiment was done in triplicate.

Somewhat surprisingly, treatment of cells with Haloperidol or (+)-PTZ did not change the interaction profile for the vast majority of proteins (Fig. 5A, B, Supplemental Tables 2 and 3). For this analysis, we considered proteins that were enriched at least two-fold (1.0 fold in the log2 scale) in the drug treated sample in comparison to the control and with a p value of at least 0.05 as being significantly changed. Using these criteria, eight proteins displayed increased binding to S1R in the presence of Haloperidol (Fig. 5A) and thirteen proteins displayed increased binding to S1R in the presence of (+)-PTZ (Fig. 5B). Interestingly, even within this small set, three proteins were shared between the two drug-treated samples, FDFT1 which encodes Squalene synthase, PCKS9 which encodes Proprotein Convertase Subtilisin/Kexin Type 9, and GARS which encodes glycyl-tRNA-synthetase. Although Haloperidol is considered an antagonist and (+)-PTZ is considered an agonist of S1R, a recent structural study found that both compounds bind within the same binding pocket of S1R (29). Thus, in addition to unique structural changes that might be induced upon ligand binding, there might also be some common protein interactions that are promoted by Haloperidol and (+)-PTZ.

**Figure 5:**
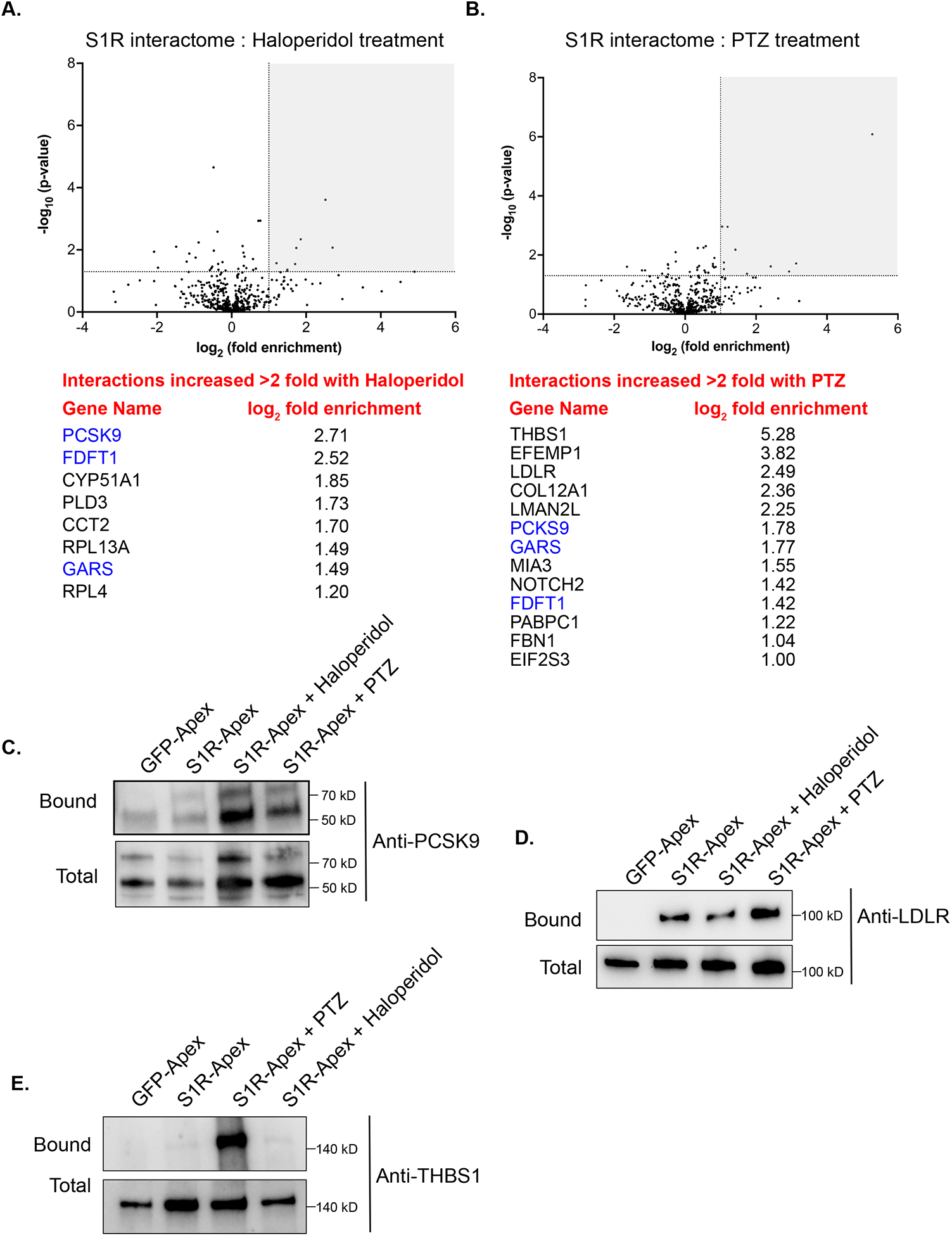
The ligand-dependent S1R interactome. **(A)** A volcano plot comparing the interactome of S1R-Apex under untreated conditions versus treatment with 25uM Haloperidol for 24 hours. A line demarcating 2-fold enrichment and a p value of 0.05 is shown. The grey shaded box indicates proteins that show at least 2-fold greater binding to S1R under Haloperidol treatment conditions with a p value of at least 0.05. A list of these proteins is shown below the volcano plot. **(B)** A volcano plot comparing the interactome of S1R-Apex under untreated conditions versus treatment with 20uM (+)-PTZ for 24 hours. The layout is similar to panel A. The proteins in blue were shown to have increased interaction with S1R under both Haloperidol and (+)-PTZ treatment conditions. **(C-E)** Biotinylated proteins were purified from HeLa cells stably expressing either GFP-Apex or S1R-Apex that were untreated or treated with either Haloperidol or (+)-PTZ. The proteins were purified using streptavidin conjugated beads. The bound proteins were eluted and analyzed by western blotting using antibodies against PCSK9 (C), LDLR (D) or Thrombospondin1 (E). Haloperidol treatment increased the interaction between S1R and PCSK9, whereas (+)-PTZ treatment increases the interaction between S1R LDLR and Thrombospondin1.

For follow-up studies, we decided to focus our efforts primarily on PCSK9 and LDLR. PCSK9 (Proprotein Convertase Subtilisin/Kexin Type 9) displays increased binding to S1R under conditions of Haloperidol treatment and to a lesser extent under PTZ treatment conditions, whereas LDLR (Low density lipoprotein receptor) binds more efficiently to S1R upon treatment with (+)-PTZ. Disease-causing variants in PCSK9 and loss of function mutations in LDLR are associated with familial hypercholesterolemia (43,44). PCSK9 is a secreted protein and extracellular PCSK9 binds to LDLR present on the cell surface. This results in endocytosis of LDLR and targeting of LDLR to the lysosome for degradation (45). Increased turnover of LDLR results in high serum cholesterol levels(46). Thus, both proteins play an important role in cholesterol metabolism. In addition, FDFT1, which encodes Squalene synthase, and CYP51A1, which encodes Lanosterol 14-alpha demethylase, are also a critical players in the cholesterol biosynthetic pathway (47,48).

Cells expressing either GFP-Apex, S1R-Apex (untreated), or S1R-Apex treated with either Haloperidol or (+)-PTZ were labeled and the binding reaction was performed as before. Bound proteins were eluted and analyzed by western blotting using an antibody against PCSK9 (Fig. 5C) or LDLR (Fig. 5D). Consistent with the proteomics results, treatment of cells with Haloperidol increased the amount of PCSK9 that was labeled and precipitated by S1R-Apex (Fig. 5C). (+)-PTZ treatment also resulted in an enrichment of PCKS9 in the S1R-Apex bound sample. Similarly, treatment with (+)-PTZ increased the amount of LDLR that was labeled and precipitated in the S1R-Apex sample (Fig. 5D). Another protein that also displayed an increased association with S1R in the presence of (+)-PTZ was Thrombospondin1 (THBS1), a finding that was also validated by western blotting (Fig. 5B, E).

During the course of these experiments, we noticed that Haloperidol treatment resulted in an increase in PCSK9 levels in the total fraction (Fig. 5C). To further test this point, we repeated the treatment in triplicate and examined cell lysates using antibodies against PCSK9 and Gapdh (as a loading control). This was indeed the case (Fig. 6A, B). A slight increase in PCSK9 levels were also noted in the (+)-PTZ treated sample. However, this increase did not reach statistical significance (Fig. 6A, B). As noted previously, PCSK9 is a secreted protein. We therefore monitored the secretion of PCSK9 under conditions of Haloperidol treatment. Culture supernatants were collected and analyzed by western blotting. In comparison to untreated and (+)-PTZ treated cells, culture supernatant from Haloperidol treated cells contained significantly more PCSK9 (Fig. 6C, D). In order to determine whether Haloperidol treatment increased global secretion we examined culture supernatants using an antibody against Thrombospondin1. In contrast to PCSK9, Thrombospondin1 was slightly decreased in culture supernatants from Haloperidol treated cells (Fig. 6C, D). These findings suggest that Haloperidol does not globally affect protein secretion and that the effect on PCSK9 is likely to be specific. Interestingly, (+)-PTZ treatment also reduced the secretion of Thrombospondin1 (Fig.6C, D).

**Figure 6:**
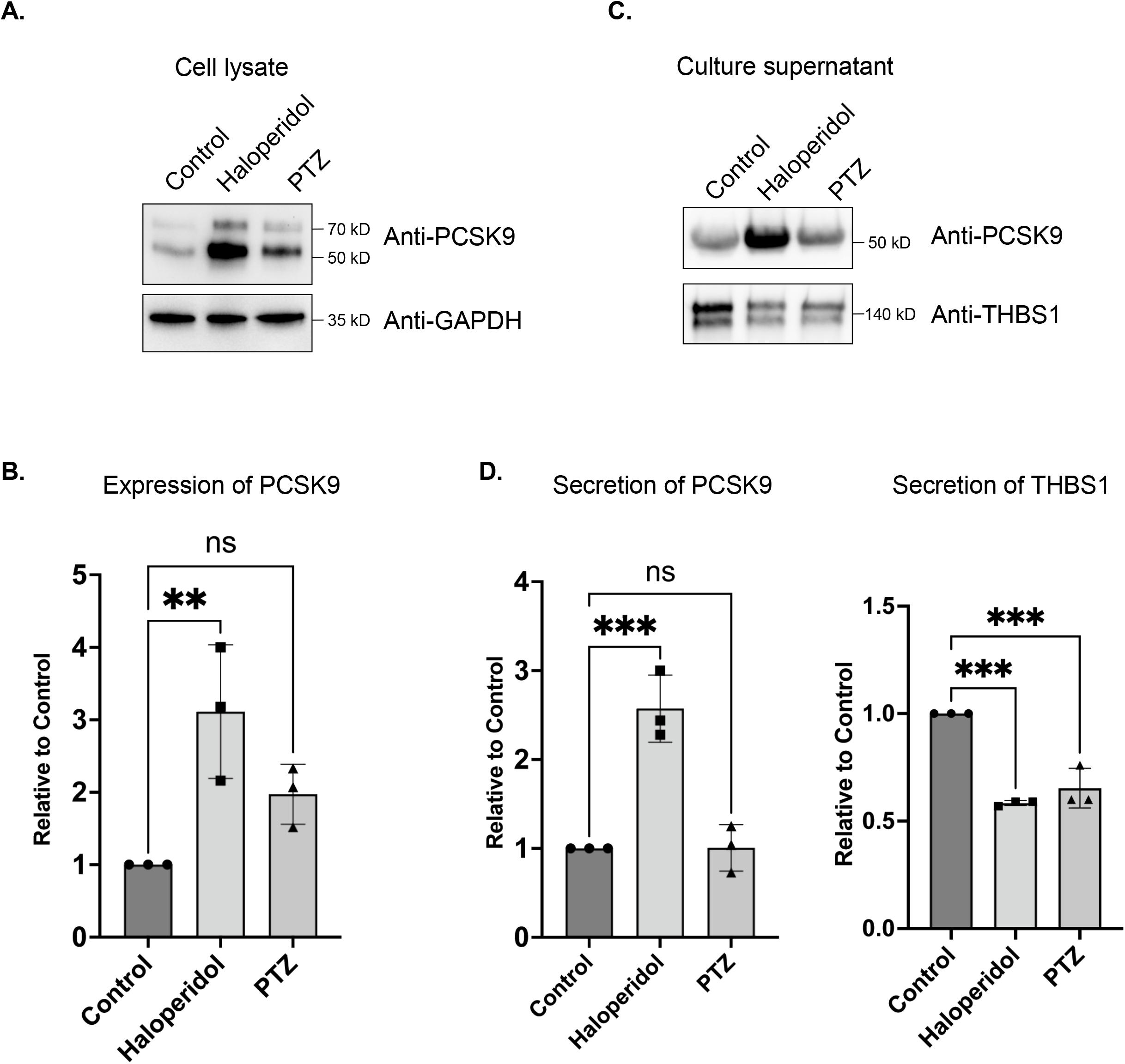
The effect of S1R ligand binding on PCSK9 and Thrombospondin1. **(A)** Lysates were prepared from HeLa cells that were either untreated or treated with either Haloperidol or (+)-PTZ. The lysates were analyzed by western blotting using antibodies against PCKS9 and GAPDH. **(B)** The above experiment was repeated in triplicate and the level of PCSK9 was quantified relative to the control untreated sample. A one-way Anova was used for these analyses; **p<0.01, ns = not significant. Haloperidol treatment results in the upregulation of PCSK9 levels. **(C)** HeLa cells were left untreated or were treated with either Haloperidol or (+)-PTZ for 24 hours. For the last four hours of this treatment, the media was replaced with serum free medium. The cells were then cultured for the final four hours. The culture supernatant was collected, concentrated and analyzed by western blotting using antibodies against either PCSK9 or Thrombospondin1. **(D)** The above experiment was repeated in triplicate and the level of secreted PCSK9 and Thrombospondin1 was quantified relative to the untreated sample. A one-way Anova was used for these analyses; ***p<0.001, **p<0.01, ns = not significant. Haloperidol treatment results in increased secretion of PCKS9. However, the sercretion of Thrombospondin1 is decreaed under these same conditions.

Given these findings, we asked whether Haloperidol treatment would affect the intracellular localization of PCSK9. Cells were treated with Haloperidol or (+)-PTZ and then processed for immunofluorescence using an antibody against PCSK9. PCSK9 was found in small cytoplasmic foci and although the intracellular level of PCSK9 was increased upon Haloperidol treatment, we did not observe any significant changes in its localization pattern (Fig. 7A-C). We next examined the localization of LDLR under these same treatment conditions. In control cells, LDLR was diffusely localized within the cytoplasm with occasional localization within small puncta (Fig. 7D). Interestingly, Haloperidol treatment caused a dramatic relocalization of LDLR to large intracellular foci (Fig. 7E). A similar, but milder phenotype was observed in (+)-PTZ treated cells (Fig. 7F). Thus, although Haloperidol and (+)-PTZ do not affect the cellular level of LDLR, both drugs affect the intracellular localization of the receptor.

**Figure 7:**
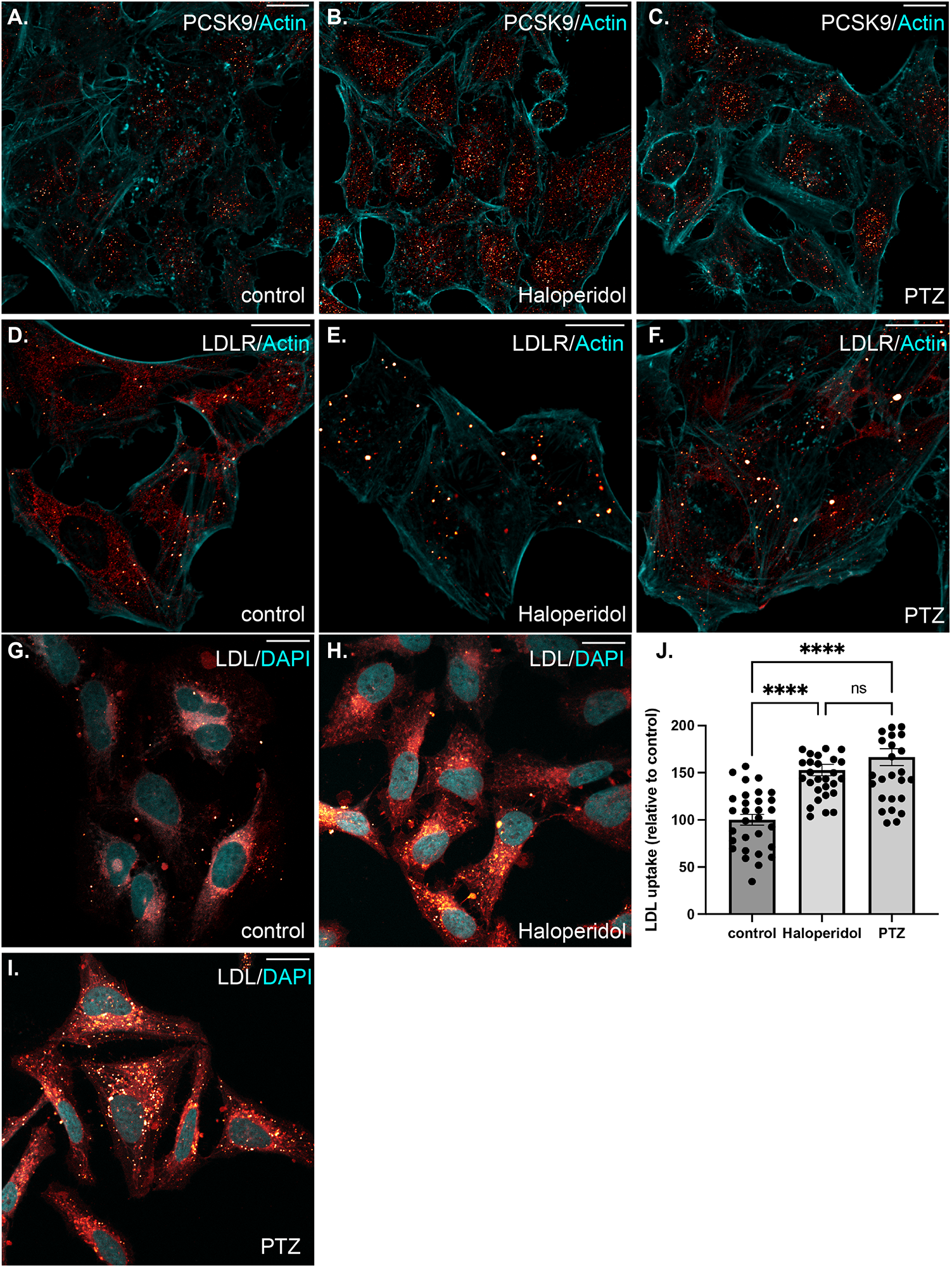
The effect of S1R ligand binding on the intracellular localization of PCSK9 and LDLR, and on LDL uptake. **(A-C)** HeLa cells were untreated (A) or treated with either Haloperidol (B) or (+)-PTZ (C) for 24 hours. The cells were then fixed and processed for immunofluorescence using an antibody against PCSK9. The cells were also counterstained using Alexa-633 conjugated Phalloidin to reveal the actin cytoskeleton (cyan). The signal for PCSK9 is depicted using a red to white range indicator, also known as a look-up-table (LUT). Low intensity pixels are shown red and high intensity pixels are shown in white. Consistent with western blotting results, the intracellular level of PCSK9 is increased in Haloperidol treated cells. **(D-F)** The cells were treated as in the above panels. The cells were then fixed and processed for immunofluorescence using an antibody against LDLR. The cells were counterstained using Alexa-633 conjugated Phalloidin (cyan). The signal for LDLR is shown using the same red to white LUT. Haloperidol treatment causes LDLR to accumulate within large intracellular puncta. A similar but milder phenotype was observed in (+)-PTZ treated cells. **(G-I)** HeLa cells were untreated (G) or treated with either Haloperidol (H) or (+)-PTZ (I) for 24 hours. The cells were then incubated with fluorescently labeled LDL. After two hours of incubation, the cells were fixed and counterstained with DAPI to reveal nuclei (cyan). The signal for LDL is shown using the red to white LUT. **(J)** The LDL assay was repeated in triplicate. The fluorescence intensity per cell was calculated for twenty frames for each treatment condition. This was used to determine the mean intensity for each treatment. The LDL uptake value for the Haloperidol and (+)-PTZ treated samples are reported relative to the control sample. A one-way Anova was used for these analyses; ****p < 0.0001, ns= not significant. The ability of cells to endocytose LDL is increased upon treatment with either Haloperidol or (+)-PTZ.

LDLR present on the cell surface binds to extracellular LDL resulting in its endocytosis. Internalized LDL is trafficked through the endocytic pathway to the lysosome. Lysosomal degradation of LDL releases cholesterol, which can then be used by the cell(49,50). We therefore examined the ability of cells that were treated with Haloperidol or (+)-PTZ to endocytose labelled LDL. In comparison to untreated cells, both (+)-PTZ and Haloperidol treatment resulted in increased LDL uptake (Fig. 7G-J). This result was somewhat surprising for Haloperidol treated cells. Higher levels of PSCK9 are generally thought to result in lower LDLR levels and therefore reduced LDL uptake. Possible scenarios that might explain this unanticipated result are discussed below.

Collectively, our findings indicate that S1R interacts with a large subset of proteins. Ligand dependent interactome changes involve proteins that are involved in cholesterol metabolism as well as proteins that are secreted into the extracellular space. The therapeutic benefit of S1R activation likely involves these and other interactome changes.

## DISCUSSION

We report here use of a cell line-based proximity labeling assay to probe the S1R interactome under native conditions and in a ligand-dependent manner. This methodology, combined with quantitative mass spectrometry-based proteomics, identified over 200 S1R-interacting proteins.

The recently published crystal structure of S1R shows that it consists of a single transmembrane segment with a short cytoplasmic tail and a large luminal ligand-binding domain(28). High resolution electron microscopy studies using an S1R-GFP-APEX2 fusion protein are consistent with luminal localization of the ligand-binding S1R C-terminus(27). Importantly, our experiments also involved tagging S1R with Apex on the C-terminus of the protein. Our results indicate that the majority of S1R interacting proteins reside within the ER lumen, the ER membrane, or within the secretory pathway. This provides functional evidence for a luminal localization of the ligand-binding domain of S1R, and is consistent with structural studies.

Previous studies indicate that S1R can influence the cellular function of many proteins(19,51). A proposed mechanism for S1R’s multiple intracellular effects has been through direct protein interactions, and S1R has been termed a ligand-operated molecular chaperone(1,3). In support of this model, experiments performed in Chinese Hamster Ovary (CHO) cells indicate that S1R forms a complex with the chaperone protein, BiP, which is known to play a central role in protein folding and quality control(1). Studies also showed that activation of S1R by the agonist (+)PTZ, led to dissociation of S1R from BiP. Release from BiP was proposed to trigger S1R’s multiple interactions with client proteins(1,52).

Our results (Fig. 2) suggest a strong interaction between S1R and BiP, consistent with previous studies. However, interestingly, we found no evidence of S1R-BiP dissociation when HeLa cells were exposed to either Haloperidol or (+)-PTZ (Fig. 5). In addition, in native condition HeLa cells (without antagonist or agonist treatment), S1R appears to be capable of interacting not only with BiP, but with hundreds of other proteins (Fig 2). Furthermore, the number of S1R-protein interactions does not substantially increase with exposure to ligands (Fig. 5). In contrast to previous studies, our results offer a comprehensive and quantitative view of the S1R interactome, and future work should address how S1R-protein interactions vary dependent on cell type analyzed, ligand specificity and exposure time as well as cell culture conditions.

Our data do show a large, robust and reproducible S1R proteomic interactome, consistent with the paradigm of S1R-mediated chaperone function. In addition, we find that S1R interacts with other known ER chaperone proteins, including Protein Disulfide Isomerase (PDI) and Calnexin. Moreover, our studies indicate a strong interaction between S1R and the ER translocation complex. We have validated the interaction between S1R and Sec61 alpha (Sec61α). The Sec61 complex forms the polypeptide-conducting channel that facilitates membrane association or translocation of nearly every newly synthesized polypeptide that is targeted to proceed through endo- or exocytic pathways(33). In addition, experiments shown in Figure 3 provide evidence that that Sec61α is functionally linked to S1R, because inhibition of protein translocation through Sec61 leads to the specific dissociation of S1R. Overall, consistent with previous work, our studies support ER-localized chaperone functionalities as central to the mechanism of S1R’s pleotropic effects.

Interestingly, our results also suggest involvement of S1R in modulation of protein secretion. Proteins destined for secretion are known to move from ER exit sites through the ER-Golgi intermediate compartment (ERGIC)(38). As shown in Figure 2B, components of the ERGIC are enriched within the S1R interactome. This finding was validated by western blot analysis of pellets using an antibody against Lman1, a marker protein of the ERGIC, and a protein that was highly enriched in the S1R interactome (Fig. 4A and Supplemental Table 1)(53,54). In addition to validating this biochemical interaction, we observed significant co-localization between S1R and GFP-Lman1 (Fig. 4B). Our results are consistent with previous studies that implicate S1R as important for regulation of the cellular secretome. For example, stimulation of S1R with (+)-PTZ leads to increased astrocytic release of brain-derived neurotrophic factor (BDNF), a neurotrophin that supports neuronal growth and survival(55). In addition, S1R has been shown to regulate levels of several other proteins that are processed and transported through the cellular secretory pathway(56,57). Future studies are needed to clarify the role of S1R in modulation of the cellular secretome.

In addition to examining the S1R interactome under native conditions, we also determined how the interactome changes when cells are treated with either the S1R antagonist, Haloperidol, or the agonist, (+)-Pentazocine. Results were notable for enrichment of proteins integral to secretion and extracellular matrix formation as well as cholesterol biosynthesis and metabolism (Fig. 5). Consistent with the latter result, previous studies have shown that S1R interacts with cholesterol and have implicated S1R in lipid metabolism and transport(58-60). However, the mechanisms that underlie these interactions are not well understood.

Our results indicate that under conditions of cellular exposure to Haloperidol, S1R interacts with the enzyme Proprotein convertase subtilisin/kexin type 9 (PCSK9). In addition, exposure to (+)-PTZ strengthens the interaction between S1R and PCSK9 and between S1R and the Low-Density Lipoprotein Receptor (LDLR) (Fig. 5). PCSK9 and LDLR are known to interact with each other and both are medically important because mutations in both genes are associated with familial hypercholesterolemia (43,61-63). LDLR, situated on cellular membranes, binds to and mediates endocytosis of cholesterol-rich LDL from the extracellular space. The endocytosed LDL is eventually broken down and cholesterol is released for use by the cell. Interaction between PCSK9 and LDLR has been shown to lead to lysosome-mediated destruction of LDLR and therefore to decreased levels of LDLR at the cellular surface(44). In general, a decrease in cell surface LDLR levels increases concentration of systemically circulating LDL-cholesterol(44). Therefore, inhibition of PCSK9 is clinically useful for treatment of hypercholesterolemia, and inhibitors of PCSK9 have been FDA-approved as cholesterol-lowering therapeutics(45).

Our findings indicate that Haloperidol treatment not only increases the S1R-PCSK9 interaction, but also leads to increased levels of intracellular and extracellular (secreted) PCSK9 (Fig.6). Somewhat surprisingly, LDLR levels were unaffected by Haloperidol treatment, at least under the conditions of our experimental setup (Fig. 5D). This finding was unexpected given the documented role of PCSK9 in mediating turnover of LDLR. However, the organ that is mostly responsible for producing and secreting PCSK9 is the liver. Secreted PCSK9 can then alter the level of LDLR in distal tissues, a scenario that is different from the cell culture setup used in our experiments. Another possibility is that treatment with Haloperidol for an extended duration might be required to observe a PCSK9-induced reduction in LDLR levels. Although Haloperidol treatment did not affect the level of LDLR, the intracellular localization of LDLR was dramatically altered. Instead of mostly diffuse cytoplasmic staining, LDLR was localized to large puncta in Haloperidol treated cells (Fig. 7). At present, the mechanism that causes this relocalization of intracellular LDLR is unknown.

In addition to exploring effects of Haloperidol, a known S1R antagonist, on the S1R interactome, we also evaluated the effects of the high affinity S1R agonist, (+)-Pentazocine, on S1R-protein interactions. Most of the interactions that are strengthened by (+)-PTZ correspond to secreted and extracellular matrix proteins such as Thrombospondin1(THBS1), Collagen12 (COL12A1) and Fibulin 1 and 3 (EFEMP1 and FBN1 respectively) (Fig. 5). For this set, we validated the interaction between S1R and Thrombospondin1 (Fig. 5E). In addition, proteins involved in cholesterol metabolism and homeostasis such as squalene synthase (FDFT1), PCKS9, and LDLR were also enriched in this dataset. Consistent with our proteomics results, (+)-PTZ treatment strengthened the interaction between S1R and LDLR (Fig. 5D). The intracellular localization of LDLR was also altered upon (+)-PTZ treatment, but to a lesser degree than when cells were treated with Haloperidol (Fig. 7).

The common theme between (+)-PTZ and Haloperidol treatment appears to involve cholesterol metabolism. Both ligands affect the localization of intracellular LDLR and treatment with both compounds promoted the cellular uptake of labeled LDL (Fig. 7). S1R ligand-dependent modulation of PCSK9 and LDLR may impact cholesterol levels and metabolism systemically or via tissue and cell type-specific effects. These findings are critically important because medications with S1R affinity are already in widespread use(13,64-66). Additional studies are needed to understand the mechanism by which S1R ligand binding affects cholesterol homeostasis and to assess the systemic ramifications of these changes.

Collectively, our findings indicate that S1R interacts with a large subset of proteins. Somewhat unexpectedly, many interactions are not significantly altered upon treating cells with either a classical antagonist or agonist. It should be noted, however, that our analysis only considered proteins showing a greater than two-fold, ligand-dependent, S1R interaction change as significant. In an organismal setting, proteins displaying more subtle interaction changes upon ligand binding might nevertheless elicit physiological effects. Furthermore, the time scale of ligand incubation might affect the degree and nature of interactome changes. Our interactome experiments were performed after 24 hours of ligand treatment. Additional time points might have revealed different interactome changes. A final consideration is that the S1R interactome might be different in different cell types and tissues. Despite these caveats, S1R agonists and antagonists are under consideration for an ever-widening spectrum of pathologies ranging from COVID-19 treatment to cancer diagnosis, chronic pain remedies, and neurodegenerative disease therapeutics(14-16,40). Our results are therefore critically relevant to a broad range of translational outcomes.

## EXPERIMENTAL PROCEDURES

### DNA constructs cell lines

The S1R-Apex construct was generated by cloning a gene synthesized product containing the cDNA sequence for mouse S1R into the pCDNA3_Sec61B-V5-APEX2 vector (Addgene plasmid 83411) (67). Gene synthesized products were obtained from Genewiz. Gibson assembly was used to replace the Sec61B cDNA in this vector with the sequence for S1R. A similar strategy was used to construct the GFP-Apex plasmid. These constructs were then subcloned into the pDONR221 Gateway vector (Life Technologies) and moved into the pAAVS1-P-CAG-DEST vector (Addgene plasmid 80490) (30). This vector enables insertion into the AAVS1 safe harbor locus present in human cell lines and also enables selection of properly integrated cells using Puromycin. All final constructs were verified by sequencing prior to use. The S1R-Apex and GFP-Apex plasmids were co-transfected into either HeLa cells (ATCC; CCL-2) or HEK293T cells (ATCC; CRL-3216) along with pXAT2 (Addgene plasmid 80494) (30). The pXAT2 vector expresses the guide RNA for the AAVS1 locus. Effectene (Qiagen) was used as the transfection reagent. Two days after transfection, stable cells were selected using 0.5ug/ml (HeLa) or 1ug/ml (HEK293T) of Puromycin (Millipore-Sigma). The plasmid expressing GFP-Lman1 was obtained from Addgene (Addgene plasmid 166942) (68).

### Immunofluorescence

Immunofluorescence was performed as previously described(69). In brief, cells were fixed in 4% formaldehyde (Pierce, ThermoFisher) for 5 min at room temperature. The cells were permeabilized by washing in PBST (PBS + 0.1% Triton X-100). The primary antibody was incubated in blocking buffer (PBS + 5% Normal goat serum) overnight at 4°C. Next, the samples were washed three times in PBST and incubated for one hour at room temperature with the fluorescent secondary antibody (Goat anti-mouse or Goat anti-rabbit antibodies conjugated with either Alexa488 or Alexa555, 1:400 dilution, Life technologies) in the same blocking buffer. The samples were then washed four times with PBST. In order to visualize nuclei, the cells were stained with DAPI. For visualizing the actin cytoskeleton Alexa633 conjugated Phalloidin was used (Life technologies; 1:400). For detecting biotinylated proteins, the samples were incubated with Streptavidin Alexa647 (Life technologies; 1:1200). The Strep674 was added to the sample at the same time as the secondary antibody. The cells were mounted onto slides using Prolong Diamond (Life technologies).

### Antibodies

The following antibodies were used: V5 was used to visualize GFP-Apex and S1R-Apex (ThermoFisher; 1:10,000 for western, 1:1000 for immunofluorescence); PDI (Cell Signaling; 1:1000 for western); Calnexin (Cell Signaling; 1:1000 for western); Sec61alpha (Santa Cruz; 1:100 for western); Sigma1 Receptor (1:500 for western) (70); Lman1 (Abcam; 1:1000 for western); PCSK9 (Abcam; 1:3000 for western, 1:300 for immunofluorescence); LDLR (Novus biologicals; 1:1000 for western, 1:100 for immunofluorescence); Thrombospondin (Abcam; 1:250 for western); Gapdh (Santa Cruz; 1:2000 for western).

### Protein purification

Biotin labeling was performed based on a previously published protocol (23). HeLa cells stably expressing either GFP-Apex or S1R-Apex were plated on 100mm culture dishes in DMEM (ThermoFisher, Waltham, MA) supplemented with 10% FBS (Atlanta Biologicals, Atlanta, GA) and 1% penicillin/streptomycin (ThermoFisher). For each replicate three 100mm dishes of cells expressing GFP-Apex and three 100mm dishes of cells expressing S1R-Apex were used. This corresponds to approximately 2.5mg of total protein from each sample. A total of three biological replicates were processed for proteomic analysis. The following day after seeding, the cells were either untreated (GFP-Apex and S1R-Apex) or treated (S1R-Apex) with 25μM Haloperidol (Tocris, Minneapolis, MN) or 20μM (+)-PTZ (Millipore-Sigma, St. Louis, MO) for 24 hours. Next, 500μM Biotin-Phenol (Millipore-Sigma) in complete medium was added to the cells for 30 minutes at 37°C. After this incubation, H_2_O_2_ (Millipore-Sigma) was added to the cell to a final concentration of 1mM. The cells were incubated for 1 minute at room temperature. This solution was then removed, and the cells were washed three times in a quench solution (10mM sodium ascorbate (Millipore-Sigma), 5mM Trolox (Millipore-Sigma) and 10mM sodium azide (Millipore-Sigma) in Dulbecco’s PBS (ThermoFisher). Next, the cells were harvested and pelleted by centrifugation. The supernatant was removed, and the cells were stored at -80°C until use. The cells were lysed in RIPA buffer (50mM Tris-Cl pH 7.5, 150mM NaCl, 1% NP40, and 1mM EDTA) containing 0.2% SDS. The lysates were centrifuged at 10,000g for 5 min at 4°C. Next, the biotinylated proteins were purified by incubating the lysates with High Capacity Streptavidin Agarose beads (Pierce, ThermoFisher) overnight at 4°C. The next day, the beads were washed three times with RIPA buffer, three times with 1%SDS, three times with RIPA buffer, three times with high salt RIPA buffer (50mM Tris-Cl pH 7.5, 1M NaCl, 1% NP40, and 1mM EDTA), three times with RIPA buffer, and finally with four PBS washes. The beads then re-suspended in PBS and processed for proteomics.

### Mass spectrometry

The beads with bound proteins were reduced with dithiothreitol, alkylated using iodoacetamide in 8M urea denaturation buffer (50mM Tris-HCl, pH 8) and digested overnight in 50mM ammonium bicarbonate using trypsin (ThermoFisher) at 37°C. Digested peptides were cleaned using a C18 spin column (Harvard Apparatus) and then lyophilized. Peptide digests were analyzed on an Orbitrap Fusion tribrid mass spectrometer (ThermoFisher) coupled with an Ultimate 3000 nano-UPLC system (ThermoFisher). Two microliters of reconstituted peptide was first trapped and washed on a Pepmap100 C18 trap (5um, 0.3×5mm) at 20ul/min using 2% acetonitrile in water (with 0.1% formic acid) for 10 minutes and then separated on a Pepman 100 RSLC C18 column (2.0 um, 75-μm × 150-mm) using a gradient of 2 to 40% acetonitrile with 0.1% formic acid over 40 min at a flow rate of 300nl/min and a column temperature of 40°C. Samples were analyzed by data-dependent acquisition in positive mode using Orbitrap MS analyzer for precursor scan at 120,000 FWHM from 300 to 1500 m/z and ion-trap MS analyzer for MS/MS scans at top speed mode (3-second cycle time). Collision-induced dissociation (CID) was used as fragmentation method.

Label-free quantification analysis was adapted from a published procedure(71). Spectra were searched using the search engine Andromeda, integrated into MaxQuant (version 1.6.15.0), against Human Uniprot/Swiss-Prot database (20,379 target sequences) (uniprot-human-swissprot-Feb2020.fasta). Methionine oxidation (+15.9949 Da), asparagine and glutamine deamidation (+0.9840 Da), and protein N-terminal acetylation (+42.0106 Da) were variable modifications (up to 5 allowed per peptide); cysteine was assigned as a fixed carbamidomethyl modification (+57.0215 Da). Only fully tryptic peptides were considered with up to 2 missed cleavages in the database search. A precursor mass tolerance of ±20 ppm was applied prior to mass accuracy calibration and ±4.5 ppm after internal MaxQuant calibration. Other search settings included a maximum peptide mass of 6,000 Da, a minimum peptide length of 6 residues, 0.05 Da tolerance for orbitrap and 0.6 Da tolerance for ion trap MS/MS scans. The false discovery rate (FDR) for peptide spectral matches, proteins, and site decoy fraction were all set to 1 percent. Quantification settings were as follows: re-quantify with a second peak finding attempt after protein identification has completed; match MS1 peaks between runs; a 0.7 min retention time match window was used after an alignment function was found with a 20-minute RT search space. Quantitation of proteins was performed using summed peptide intensities given by MaxQuant. The quantitation method only considered razor plus unique peptides for protein level quantitation.

### Western blotting

For the validation experiments using western blotting, the same procedure was followed using ¼ the number of cells. 10ul of whole cell lysate was collected prior to the binding to run as the total fraction. The same binding and wash steps were used. The bound proteins were eluted by boiling in Laemmli gel loading buffer. Proteins were separated by electrophoresis on a 4-15% SDS-polyacrylamide gel (BioRad) and then transferred to a nitrocellulose membrane (ThermoFisher). The membrane was blocked with 5% nonfat milk in Tris-buffered saline-0.05% Tween 20 (TBST) for 1h at room temperature, then incubated overnight at 4°C with primary antibodies. After three washes in TBST, the membrane was incubated for 1h using an appropriate Horseradish Peroxidase (HRP)-conjugated secondary antibody (Santa Cruz Biotechnology) at room temperature. Proteins were visualized by incubating with a SuperSignal West Pico chemiluminescent substrate (ThermoFisher). A BioRad ChemidocXP was used to visualize and quantify western blot signal.

### Microscopy

Images were captured on either a Zeiss LSM 780 inverted confocal microscope or an inverted Leica Stellaris confocal microscope located within the Augusta University Cell and Tissue imaging core.

### LDL uptake assay

HeLa cells were seeded onto glass coverslips. After the cells adhered, they were either left untreated or treated with 25μM Haloperidol or 20μM PTZ for a total of 24 hours. The LDL uptake assay (Image-iT Low Density Lipoprotein Uptake kit, ThermoFisher) was performed as per the manufacturer’s instructions. Briefly, for the last 18 hours of drug treatment, the medium was replaced with serum free DMEM containing 0.3% BSA, and then incubated with 10ug/ml of labeled LDL at 37°C for 2 hours. The cells were then washed twice with PBS and fixed with 4% formaldehyde (Pierce) for 5 min at room temperature. The cells were counterstained with DAPI. The coverslips were mounted onto slides using Fluoroshield (Millipore-Sigma). Cells were imaged for quantification using a Zeiss Axio Imager D2 microscope equipped with Zeiss Zen23pro software and a high-resolution camera. For high resolution images, cells were imaged on a Zeiss 780 inverted confocal microscope.

### Software

Proteins levels were quantified using the ImageLab software (BioRad). Images were processed for presentation using Fiji, Adobe Photoshop, and Adobe Illustrator. Graphs and volcano plots were assembled using Graphpad Prism9. Statistical analysis were also performed using Graphpad Prism.

## Figure Legends

**Table 1:** A list of the top 60 S1R interacting proteins.

**Supplemental figure 1:**

HeLa cells stably expressing GFP-Apex were fixed and processed for immunofluorescence using a V5 antibody (green). The cells were also incubated with Streptavidin-647 (red) to reveal the localization of biotinylated proteins and were counterstained using DAPI (cyan). Biotinyalted proteins are observed in the nucleus and cytoplasm.

**Supplemental table 1:** List of proteins that specifically interact with S1R-Apex.

**Supplemental table 2:** List of proteins that change their association with S1R-Apex upon treatment with Haloperidol.

**Supplemental table 3:** List of proteins that change their association with S1R-Apex upon treatment with (+)-PTZ.

## Notes

### Competing Interest Statement

The authors have declared no competing interest.

