## Supplementary figures and images for "Defining the Ligand-dependent Interactome of the Sigma 1 Receptor"

### Supplemental Figure1

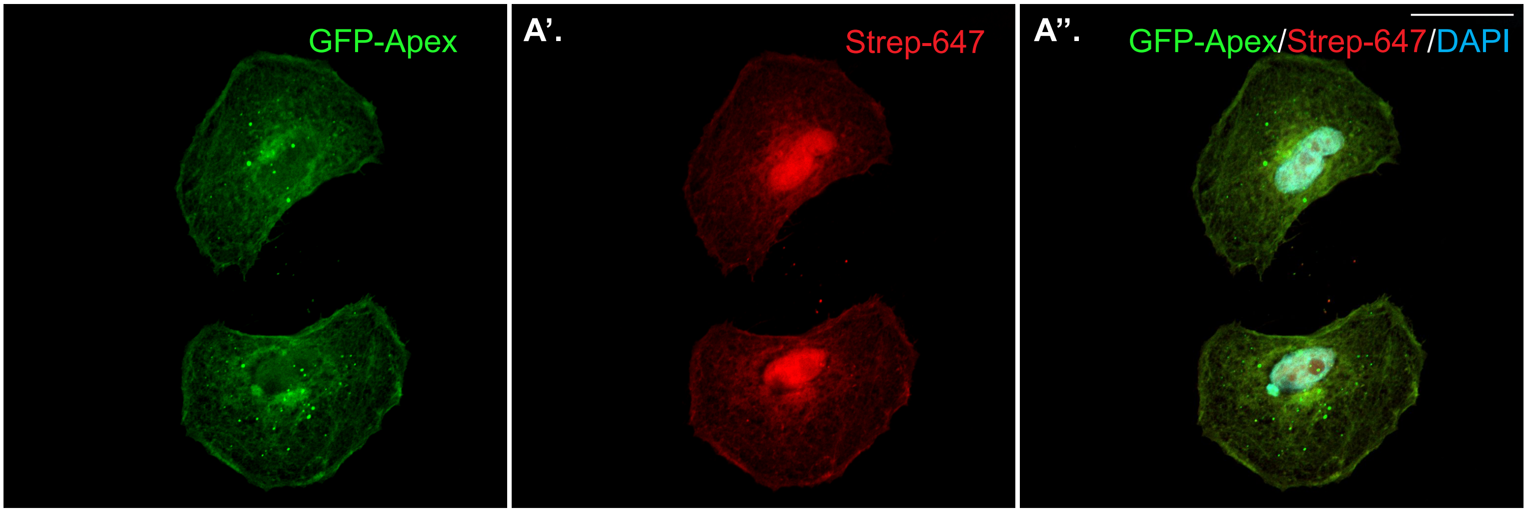


Supplemental Figure 1
